# Ultra-flexible and Stretchable Intrafascicular Peripheral Nerve Recording Device with Axon-dimension, Cuff-less Microneedle Electrode Array

**DOI:** 10.1101/2022.01.19.476928

**Authors:** Dongxiao Yan, Ahmad A. Jiman, Elizabeth C. Bottorff, Paras R. Patel, Dilara Meli, Elissa J. Welle, David C. Ratze, Leif A. Havton, Cynthia A. Chestek, Stephen W.P. Kemp, Tim M. Bruns, Euisik Yoon, John Seymour

**Affiliations:** Department of Electrical Engineering and Computer Science, University of Michigan, Ann Arbor, MI; Department of Electrical and Computer Engineering, King Abdulaziz University, Jeddah, Saudi Arabia; Department of Biomedical Engineering, University of Michigan, Ann Arbor, MI; Department of Materials Science and Engineering, Northwestern University, Evanston, IL; Departments of Neurology and Neuroscience, Icahn School of Medicine at Mount Sinai, New York, NY; James J Peters Veterans Affairs Medical Center, Bronx, NY; Section of Plastic Surgery, University of Michigan, Ann Arbor, MI; Center for Nanomedicine, Institute for Basic Science (IBS) and Graduate Program of Nano Biomedical Engineering (Nano BME), Advanced Science Institute, Yonsei University, Seoul, South Korea; Department of Neurosurgery, UTHealth, Houston, TX; Department of Electrical and Computer Engineering, Rice University, Houston, TX

## Abstract

Peripheral nerve mapping tools with higher spatial resolution are needed to advance systems neuroscience, and potentially provide a closed-loop biomarker in neuromodulation applications. Two critical challenges of microscale neural interfaces are (i) how to apply them to small peripheral nerves, and (ii) how to minimize chronic reactivity. We developed a flexible *mi*croneedle *n*erve *a*rray (MINA), which is the first high-density penetrating electrode array made with axon-sized silicon microneedles embedded in low-modulus thin silicone. We present the design, fabrication, acute recording, and chronic reactivity to an implanted MINA. Distinctive units were identified in the rat peroneal nerve. We also demonstrate a long-term, cuff-free, and suture-free fixation manner using rose bengal as a light-activated adhesive for two timepoints. The tissue response at 1-week included a sham (N=5) and MINA-implanted (N=5) group, and the response at 6-week also included a sham (N=3) and MINA-implanted (N=4) group. These conditions were quantified in the left vagus nerve of rats using histomorphometry. Micro-CT was added to visualize and quantify tissue encapsulation around the implant. MINA demonstrated a reduction in encapsulation thickness over previously quantified interfascicular methods. Future challenges include techniques for precise insertion of the microneedle electrodes and demonstrating long-term recording.

## 1. Introduction

Current implants for directly interfacing with peripheral nerves can be categorized into extra-fascicular and intrafascicular devices. Extrafascicular devices, usually conventional cuff electrodes^[1,2]^ or advanced derivations thereof^[3]^, have been proven to be effective for a large variety of electrophysiology studies and neural modulation applications^[4,5]^. The development of intrafascicular devices enabled researchers to detect and modulate neural activities at a higher resolution. To date, a variety of intrafascicular recording and/or stimulation methods have been achieved using thin-film polymer devices^[6,7]^, carbon fiber electrode arrays^[8,9]^, highly conductive carbon-based threads with low electrochemical impedance^[10–12]^, Utah arrays^[13,14]^, flexible microneedle electrode arrays^[15–17]^ and regenerative devices ^[18]^. With implantation of such devices, researchers were able to map and decode motor and/or sensory somatic or autonomic systems at a sub-fascicular level^[19,20]^.

For all types of intrafascicular devices, minimizing acute implantation damage and chronic foreign body reaction is essential. Intuitively, the difficulty of developing such devices and the corresponding implantation strategy increases when targeting nerves with small sizes such as autonomic nerves. The electrodes and supporting substrate should ideally be minimized. Furthermore, the mechanical properties of the substrate should be made from low-modulus materials to reduce the mechanical mismatch. As an example, the high-density Utah array which achieved success in sciatic nerve recordings^[21]^, is not likely a useful option for autonomic nerves due to the dimension and rigidness of the monolithic structure. Lastly, the implantation procedures should ideally be designed and performed in a way that minimizes both acute and chronic tissue damage. A challenge with the longitudinal intrafascicular electrode (LIFE) and transverse intrafascicular multi-electrode (TIME) array implantation procedure is threading the polyimide body (thin film) through the nerve with a shuttle, needle, or suture. The acute damage from insertion may results in structural and vascular damage, local compression, and tensile strain of the nerve. This method is technically challenging and may be overwhelming for small size nerves.

Another determining factor of the recording quality is the amount of tissue encapsulation over the electrodes during chronic implantation. Wurth et al.^[22]^ provide an insightful and detailed characterization of a flexible TIME array in the sciatic nerve resulting in an encapsulation thickness of 60 microns and demyelination ranging from 90 to 170 microns. The magnitude of the reactivity may be due to the acute structural damage, the chronic aggravation of even a mildly stiff structure, or simply a foreign body response to the wide polyimide substrate. There is some evidence in the central nervous system that smaller structures, even moderately rigid structures, may reduce the reactivity to an implant by reducing the size of the implant itself, such as with a sub-cellular mesh structure^[23,24]^ or a subcellular fiber^[25,26]^. The working hypothesis of many neurotechnologists is that as the electrode approaches biomimetic dimensions, the longitudinal fidelity and density will improve^[27]^.

Lastly, nerve compression induced by the cuff structure is known to cause chronic nerve damage^[28,29]^. To potentially minimize compression over the nerves, Liu. et al. developed a stretchable cuff electrode with low-modulus materials, although with dimensions better suited for large somatic nerves^[1]^.

To address the challenges discussed above and create a high spatial resolution and low tissue reactivity intrafascicular interface, we developed a silicon-based high-density microneedle penetration electrode array having axon-dimension needles. Each electrode had an independent mechanical coupling to a low-modulus substrate. Materials with good biocompatibility and stability *in vivo* such as thermal oxide^[30]^ and Parylene C were carefully selected and processed as the insulation materials. Additionally, we eliminated the cuffing or wrapping of the device to the nerve using a cuff-less photochemical tissue bond.

## 2. Results and Discussion

Microneedle nerve array (MINA) was developed to record intrafascicular neural signals from small peripheral nerves (e.g., < 1 mm in diameter) with a microneedle penetrating electrode array (Figure 1).

**Figure 1.**
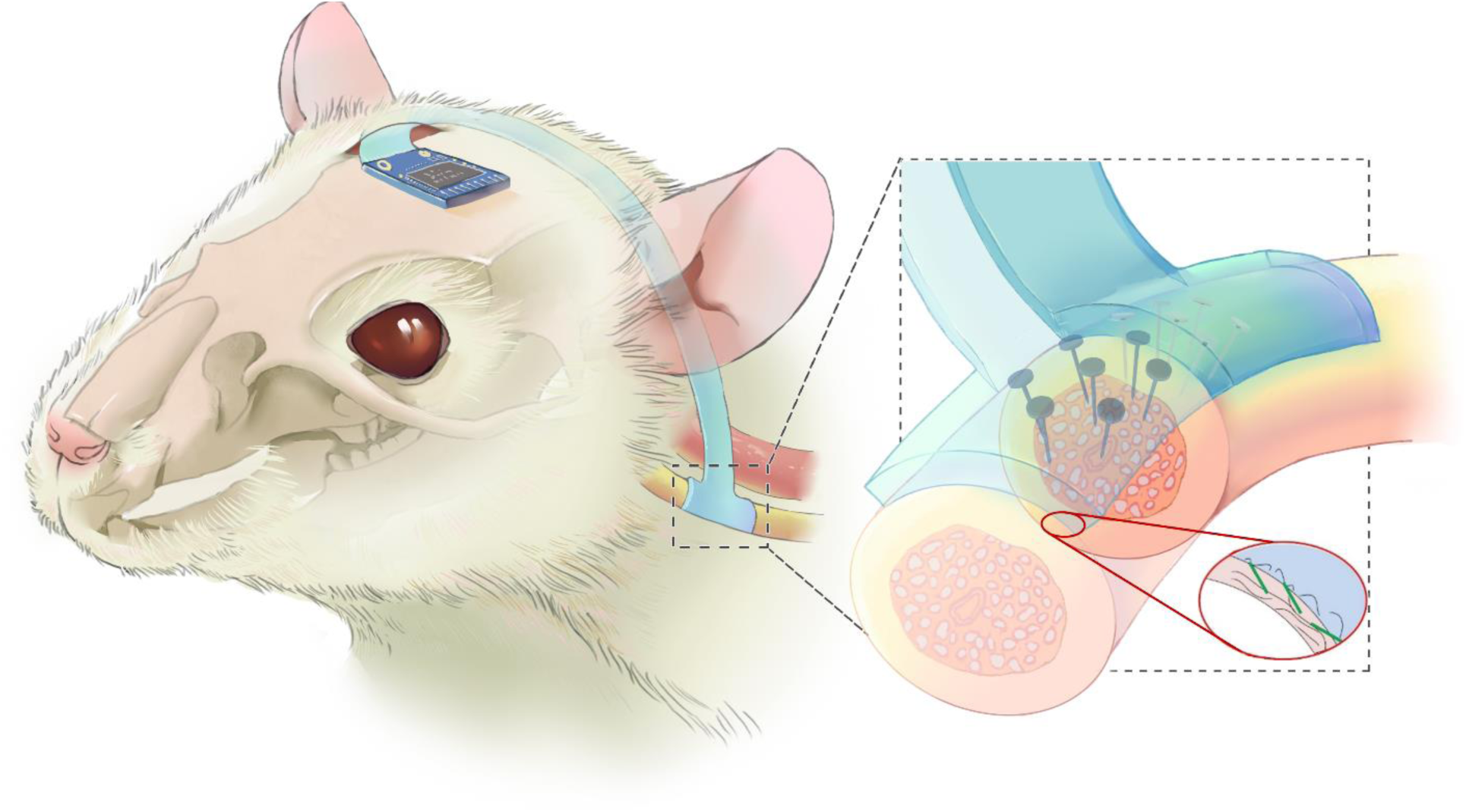
Conceptual diagram of a chronically implanted MINA in a rodent autonomic nerve. Cuff-less implantation of MINA can be achieved by bonding the substrate onto the epineurium via photochemical reaction with the axon-dimension electrodes penetrating the perineurium as demonstrated.

### 2.1 MINA design and fabrication

MINA devices were fabricated using microelectromechanical system (MEMS) fabrication technologies as further details are described in the *Experimental Section*. Shown in figure 2a, a double-sided process was developed to create MINA devices from a silicon on insulator (SOI) wafer. Boron-doped silicon wafer (ρ = 0.001-0.005 Ω·m) was chosen for its high conductivity and biostability *in vivo*^[31]^. First, we used the device layer to fabricate the conductive silicon microneedle array with a series of customized deep reactive ion etching (DRIE) steps (Figure 2 b-d) to create a sharp tip and several angles on the silicon microneedles, which we further describe in Figure S1. The microneedle array was embedded partially in polydimethylsiloxane (PDMS). A 400 nm thick Parylene C layer followed by a thick photoresist layer was coated on top leaving a ∼20 μm tall microneedle tip exposed. Upon removal of the dielectric materials over the tip, the electrode metal (Ti/Pt) was coated via sputtering. We etched the handling layer silicon to expose the backend of the microneedles. Finally, metal interconnection traces and insulating materials were patterned from the backside. The as-described combination of microfabrication processes comprising RIE microneedle shaping, dielectric layer formation and electrode metallization, can create high-density arrays of electrodes with many custom geometries (Figure S1).

**Figure 2.**
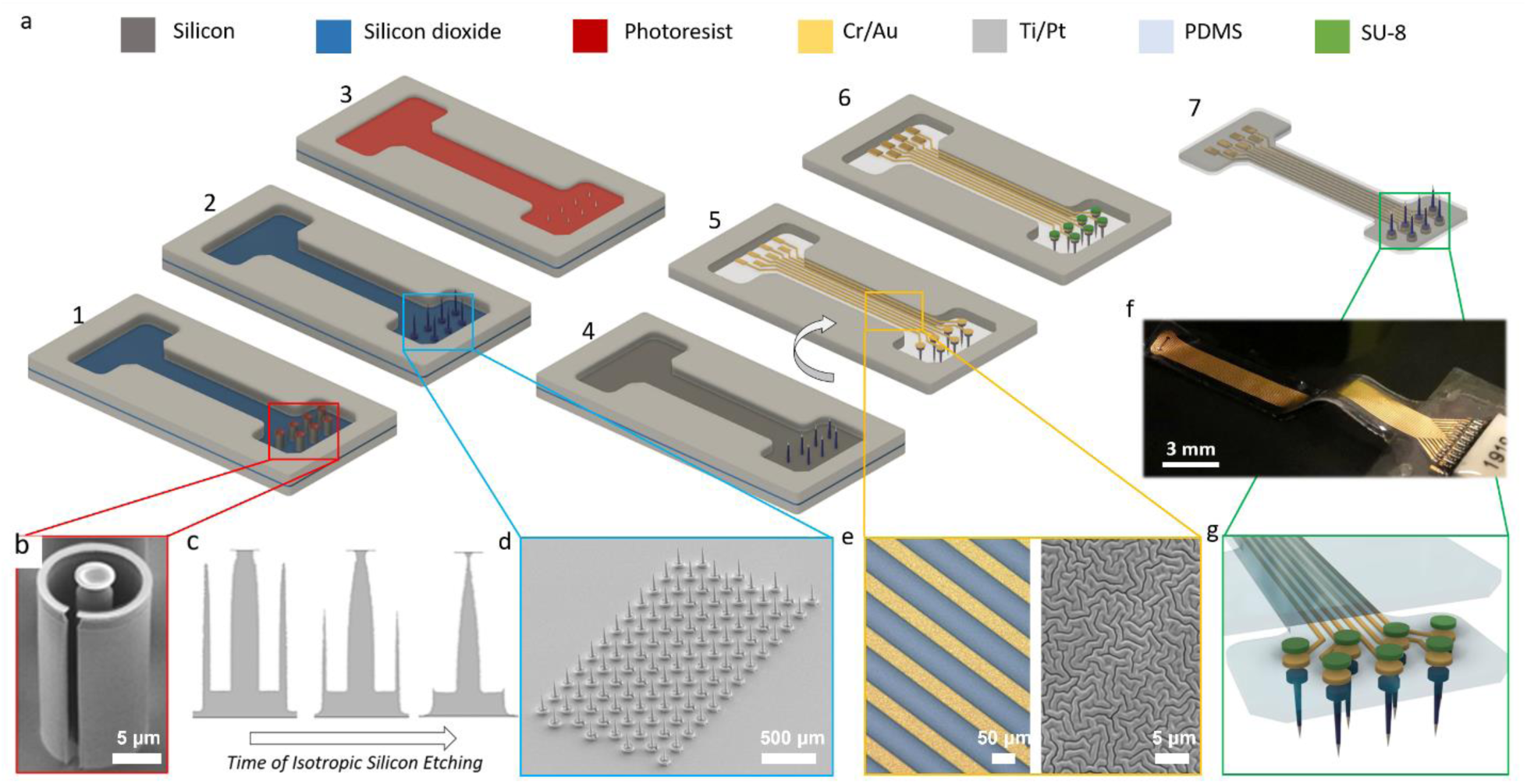
Overview of MINA microfabrication. **a)**. The major steps 1-7 of MINA fabrication. The Parylene-C coatings are not shown in this diagram for simplification. **b)**. Silicon microneedle shaping by deep reactive ion etching (anisotropic). **c)**. Microneedle shape was tapered by isotropic removal of a sacrificial ring structure. **d)**. SEM image of an array of 100 needles in a honeycomb pattern and 150-μm pitch. Thermal oxide insulation is grown on the needle. **e)**. Optical and SEM image of the thin-film metal interconnect traces with micro-wrinkle morphology for enhanced strain compliance. **f)**. Photo of a MINA device. **g)**. A zoomed-in diagram of the contact metal and protective SU-8 over the back of the microneedle electrodes.

Compared with other common dielectric coating approaches where a thin layer of organic or inorganic insulator was deposited via vapor deposition, high-quality silicon dioxide formed using thermal oxidation was shown to have good biocompatibility and longevity in the physiological environment^[30–34]^. This is the first instance of thermal oxide grown on microneedles, to our knowledge. The 70 μm thick PDMS as the substrate material enhanced the flexibility of the device and enabled the electrode array to conformally attach to the small diameter autonomic nerve. The Young’s modulus of PDMS, which is typically below 3 MPa, is much closer to the tensile Young’s modulus of nerve^[35]^ compared with other commonly used polymer materials for flexible devices.

*In vitro* characterization of the microneedle electrodes was performed in phosphate buffered saline (PBS) solution at room temperature. The impedance between each single electrode of a 16-channel MINA device and an Ag/AgCl counter electrode was measured using an LCR meter (Keysight E4980AL). Shown in Figure 3c, the average impedance at 1 kHz was 215 ± 89 kΩ measured of 3 devices with 16 electrodes each.

**Figure 3.**
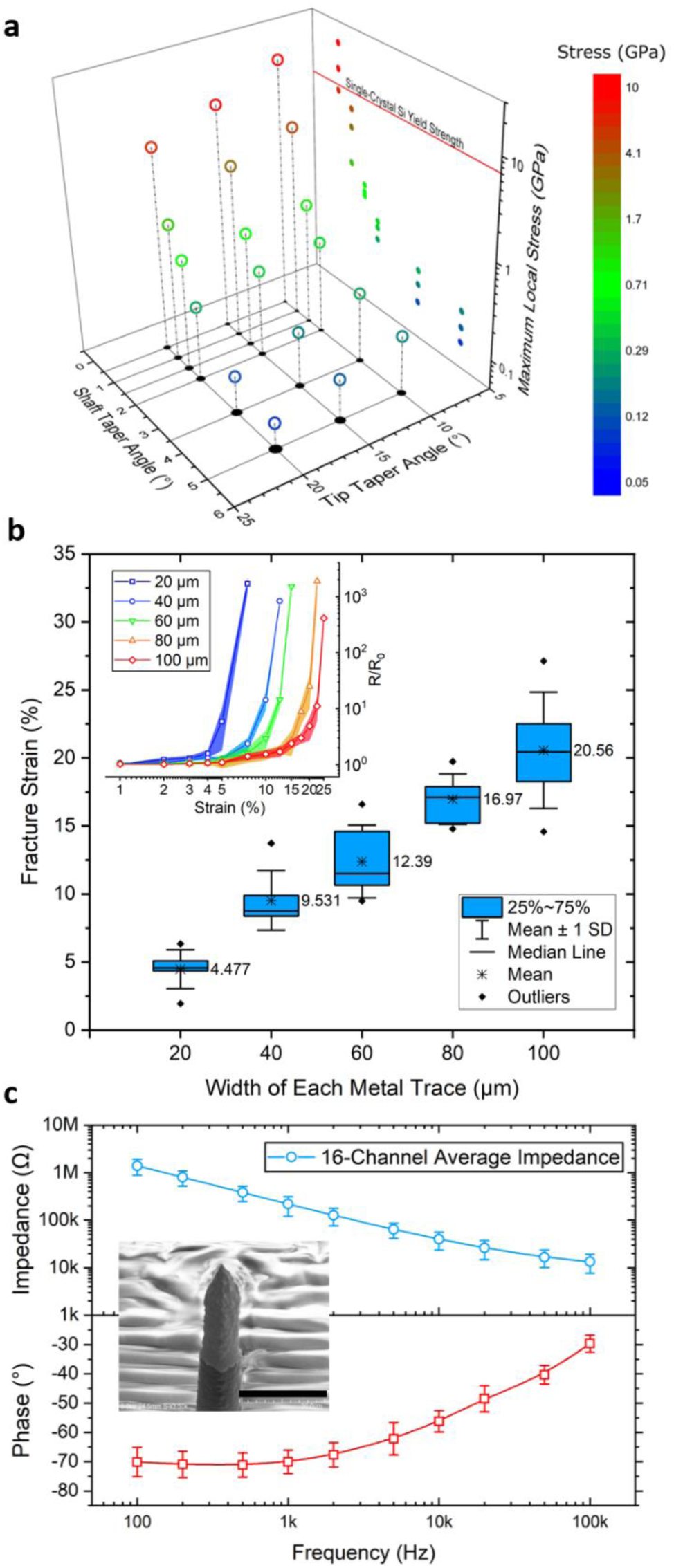
Mechanical and electrical characteristics of MINA. **a)**. COMSOL finite element method simulation of the maximum stress along the microneedle structure when 1-mN force was applied horizontally. The plot shows the maximum stress among different dual-taper designs. **b)**. Strain compliance of the thin-film micro-wrinkled metal traces of different widths. The inset shows the resistance change over strain for each width value. **c)**. Mean and standard deviation of the electrode impedance and phase measured in saline. The inset shows the Ti/Pt coated microneedle tip under SEM. The scale bar is 20 μm.

### 2.2 Mechanically robust microneedle electrode and interconnection design

We successfully fabricated microneedle electrodes with an average (along the microneedle shaft) diameter ranging from 11 to 15 μm and a sharpened tip that was thinner than 1 μm. These diameter dimensions are similar in size to myelinated somatic axons^[36]^ and just above the size of myelinated autonomic axons^[37]^, demonstrating the axon-sized nature of these microneedle electrodes. A finite element method simulation using COMSOL was performed to predict the stress distribution along the microneedle electrode. In the model, a lateral force was applied at the tip of the electrode to simulate potential distortion during insertion. According to the simulation, a “dual-taper” geometry allows the stress to be distributed relatively evenly along the microneedle. It prevents stress from localizing at either the tip or the foot of the microneedle which would create mechanical weak points. A comparison between different geometries via simulation was presented in a previous publication^[15]^. To further determine a target geometry for the fabrication, we tested different combinations of the two tapered angles. An arbitrary range of the average diameter of interest was set to be from 7.5 μm to 20 μm. Any combinations of the two taper angles which produced an average diameter within this range were modeled. Finally, taking the process capability, structure minimization, and the predicted maximum stress into consideration, we targeted a microneedle electrode design with approximately a 15° tip taper and a 1.5° shaft taper. The maximum stress along the body of the microneedle electrode with such design shown by the modeling was 1.84 GPa.

The mechanical property mismatch among the PDMS substrate, thin-film metal interconnection, and silicon microneedles can cause an open circuit especially at the edges of the metal to silicon contact. To address this issue, we applied an extra layer of SU-8 with a thickness of 2 μm on top of the metal patterns. The addition of this mechanical reinforcing layer noticeably improved the yield and robustness of MINA devices. However, although being reduced, strain over the metal interconnections is still inevitable during chronic studies with freely moving animals. We deposited and patterned the metal interconnection on a pre-strained wrinkled surface. The wrinkled surface topology was created by intrinsic stress of the Parylene C chemical vapor deposition on top of PDMS. As a result, the thin-film metal traces had uniform, wrinkled surface topology (Figure 2 e) and exhibited a high degree of strain tolerance which is a function of the trace width (Figure 3 b). Strain compliance of the as-prepared metal traces was measured using separated test samples (N=6) without embedded microneedle electrodes. The average MINA devices were fabricated with 80-μm wide metal traces.

Based on the observation during the acute recording experiments, microneedles with an average diameter between 13-15 μm (measured near the microneedle center) had a much higher insertion yield of over the microneedle electrodes with an average diameter of 11 μm or less. During chronic implantation of the un-tethered MINA devices (N=10) implanted for both 1 and 6 weeks, 11 out of 240 (4.6%) of the total microneedle electrodes were broken during or before implantation. The insertion yield results validated the mechanical robustness enabled by the “dual-taper” design when the average cross-section diameter was greater than 13 μm for the length and taper described.

### 2.3 Acute neural recordings responding to cutaneous brushing stimulation

Our primary goal in these experiments was to obtain intraneural recordings driven by controlled stimuli. Thus, we targeted the somatic peroneal nerve, which is easily accessible and has large, myelinated axons that transduce cutaneous and proprioceptive sensory inputs, instead of an autonomic nerve like the vagus nerve. As detailed in the *Experimental Section*, MINA devices were attached to the bottom of a vacuum suction tool and implanted into a peroneal nerve using a pen camera for visualization. The results shown in this section and Figure 4 were collected from one successful MINA insertion out of four trials. The pen-camera view during MINA implantation is shown in Figure 4c together with a zoom-in photo of the MINA electrode array in Figure 4d. We used similar surgical setups during the acute MINA recording trials and the chronic non-tethered MINA implantation trials (Figure S4e). Electrochemical impedance of the electrodes was measured in saline and after insertion. The channels which showed similar impedance values after insertion were considered as inserted successfully. The average impedance measured at 1 kHz after insertion was as 342 ± 46.4 kΩ at 1 kHz (N=12 out of a total of 16 channels).

**Figure 4.**
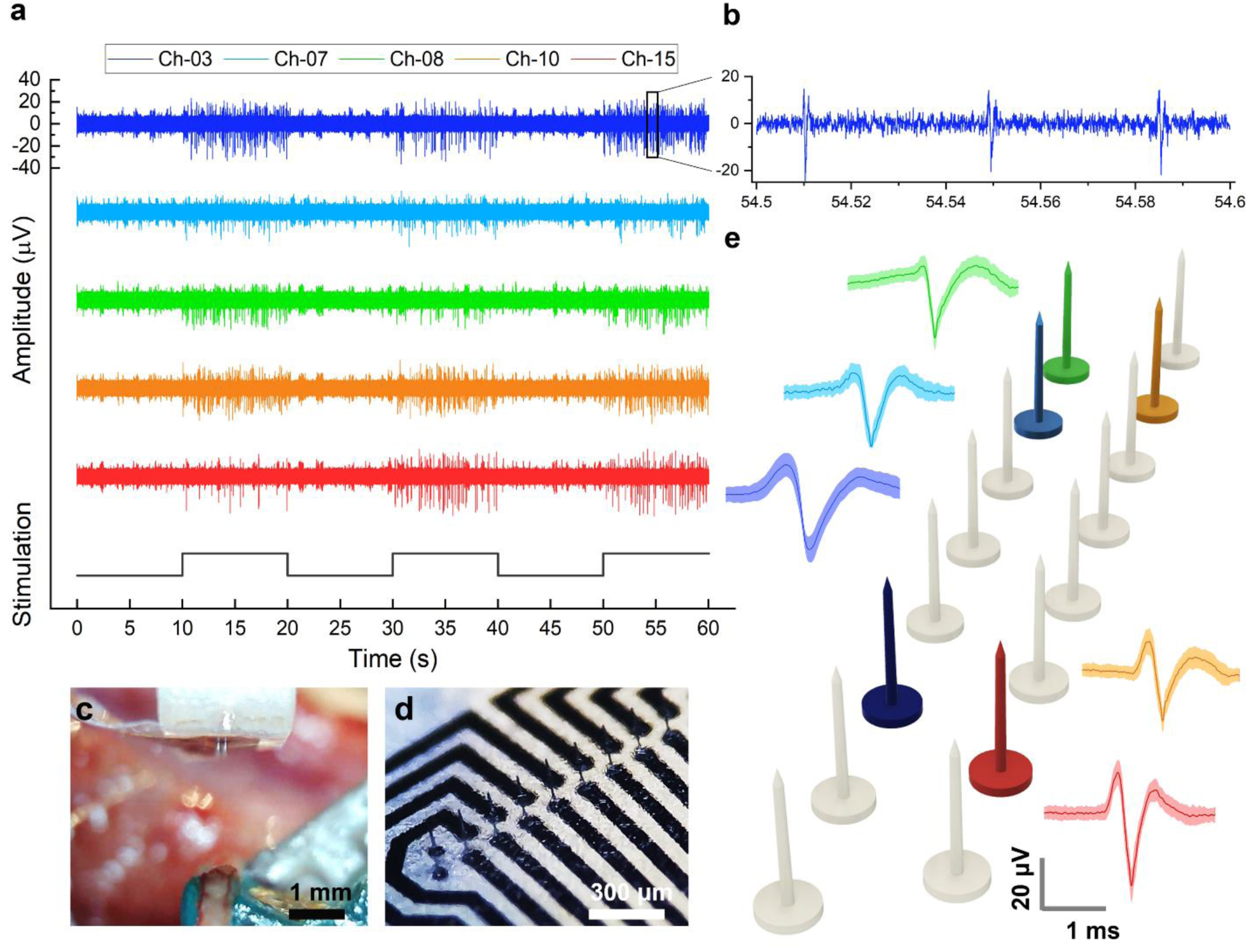
MINA intrafascicular recording from rodent peroneal nerve. **a)**. Neural signals recorded on multiple MINA channels responding to cutaneous brushing stimulation on the hind leg. **b)**. Zoom-in of one channel. **c)**. The pen camera view of the microneedle electrodes during implantation into the peroneal nerve. **d)**. Zoom-in photo of the MINA electrode array. **e)**. Example sorted spikes recorded using MINA (darker line: mean waveform; shading: standard deviation range across waveforms).

Neural recordings responding to cutaneous brushing stimulation were identified on multiple channels, with a peak-to-peak amplitude of 23 to 40 μV and an average signal-to-noise ratio of 9.74 dB. Signals from 5 representative channels are shown in Figure 4a and b. Mean and standard deviations of the waveforms of several sorted units from those channels are shown in Figure 4e, in which slight differences between channel unit waveforms can be seen. Among the 16 recording channels from the device implanted in this trial, one unit and two units per channel were identified on 9 channels and 2 channels, respectively. The channels failed to pick up spikes had higher impedance of over 2 MΩ measured after insertion. This could be caused by the electrodes landing in the high-impedance tissue rather than endoneurium or breaking of the microneedle body. We are confident that the sorted spikes are action potential recordings based on the waveform shape and width (time scale). However, we were unable to confirm if all spikes on a given channel were from a single or multiple axons due to similarity in spike waveform shapes and normal variations in amplitude. The observation of one unit typically sorted per electrode in a nerve fascicle is consistent with prior studies of microelectrode interfacing with peripheral nerves^[8,38]^. Intraneural recordings from peripheral nerves, such as shown here, may have significant utility in studying organ and limb neurophysiology and in providing sensory feedback for neuroprostheses. For example, vagus nerve recordings^[8]^ may be used to initiate neuromodulation for hypertension or gastric motility^[39]^. As MINA can be created in a wide range of channel layouts, including in a single column, it offers a new, scalable interface for recording action potentials from nearly any size of peripheral nerves.

Inserting the MINA into the somatic peroneal nerve was challenging. The most significant factors that negatively influenced the trial success rate were the small dimension of the array and the nerve, fluid in the cavity, movement of the nerve when applying force, and the difficulty of positioning a camera to visualize alignment and insertion (Figure 4c). During all the trials, microneedle electrodes fracture was observed caused by pressing them onto the rigid nerve holder instead of the nerve tissue, which was an alignment error. For single and multi-unit recording in peripheral nerves, microelectrode sites need to be in close proximity of an axon node due to the small extracellular fields^[40]^. Electrodes not penetrating the perineurium layer could be another cause of failing to record the small amplitude spike signals. An improvement of the implantation procedures and surgical tools together with a longer microneedle length may contribute to a higher trial success rate.

### 2.4 Cuff-less chronic implantation of MINA by photochemical bonding using rose bengal

The cuff-less implantation strategy developed in this work utilized photochemical reaction to create chemical bonds between the functionalized surface of the device and the epineurium layer. Among the animals implanted with untethered MINA chronically using this approach, 6 out of 8 untethered MINA devices stayed fully attached to the nerve after 1-week implantation and 4 out of 7 after 6-week implantation. The success rate may be further improved with better surgical procedures. Analysis of the chronic implantation outcome is described in the next two sections.

Upon irradiation with green light, rose bengal (RB) is excited into a primary excited state, and then decays into a triplet excited state, followed by an energy transfer to oxygen to create singlet oxygen^[44]^. This highly reactive species then initiates protein cross-linking. The bonding strength was further characterized with an *in vitro* setup. Measured force vs. displacement curves were characteristic in an almost linear increase followed by an abrupt decline of the force required denoting detachment of the sample from the nerve (Figure S2). Temperature measurements in rat, porcine, and ovine nerves were performed to evaluate laser heating (Figure S3) and the device-to-tissue bonding strength (Figure S2c). Small nerves heated quickly and therefore we were careful to limit the irradiance and time of the exposure. Tensile and shear adhesion strength by RB induced photochemical bonding were characterized at 7.12 ± 4.71 kPa (N=10) and 18.2 ± 13.4 kPa (N=10), respectively. Details of the comparison and the testing setup are provided in Table S1 and Figure S2. Some common alternatives to such an approach are using cyanoacrylate or fibrin glue. There is potential for polydopamine^[41]^ and UVA-riboflavin^[42]^ as alternative adhesives as well, due to their excellent coating abilities and biocompatibility. It is critical that the adhesive can be applied to a medical device prior to application as a thin coating to control the contact area, reduce the adhesion thickness on the device, maintain high flexibility, and can be applied on wet tissue or even in an aqueous environment^[43]^. Rose bengal also enabled precise timing of adhesion over a precise area.

During the *in vivo* implantation using this approach (Figure 4, S4), a lower power output was used which delivered the same amount of total energy but over a longer period. After inserting the microneedle array, a 532-nm laser irradiation (85 mW, 0.8 mm beam-diameter, Civil Laser) was applied through a 3D-printed diffusion lens over the back of the array (Figure S4c, h), which was the same lensing system used in our benchtop evaluations (Figure S3). The lens diffused the laser power to ∼1 W/cm^2^ over the entire area of an untethered MINA. Figure 5a shows an example of an untethered MINA bonded onto the vagus nerve after insertion. We measured the average nerve temperature increases with an infrared sensor at the time of implantation as 1.5 ± 1.6 °C in 1-week MINA surgeries and 0.7 ± 0.7 °C in 6-week MINA surgeries from 6 and 8 trials respectively. The nerve was also suspended on the nerve hook and below body temperature, therefore the absolute temperature of all measurements (26-35 °C) were within the functional range for peripheral nerves (24-37 °C)^[49]^. Across the MINA implant procedure for 1-week and 6-week rats, we measured a nerve strain of 5.9 ± 2.5% due to handling and elevating the nerve from a total of 7 trials. While this amount is within a tolerable limit for peripheral nerves (∼12%)^[50]^, it may have compromised the structure of the nerve and reduced our attachment success in some cases.

**Figure 5.**
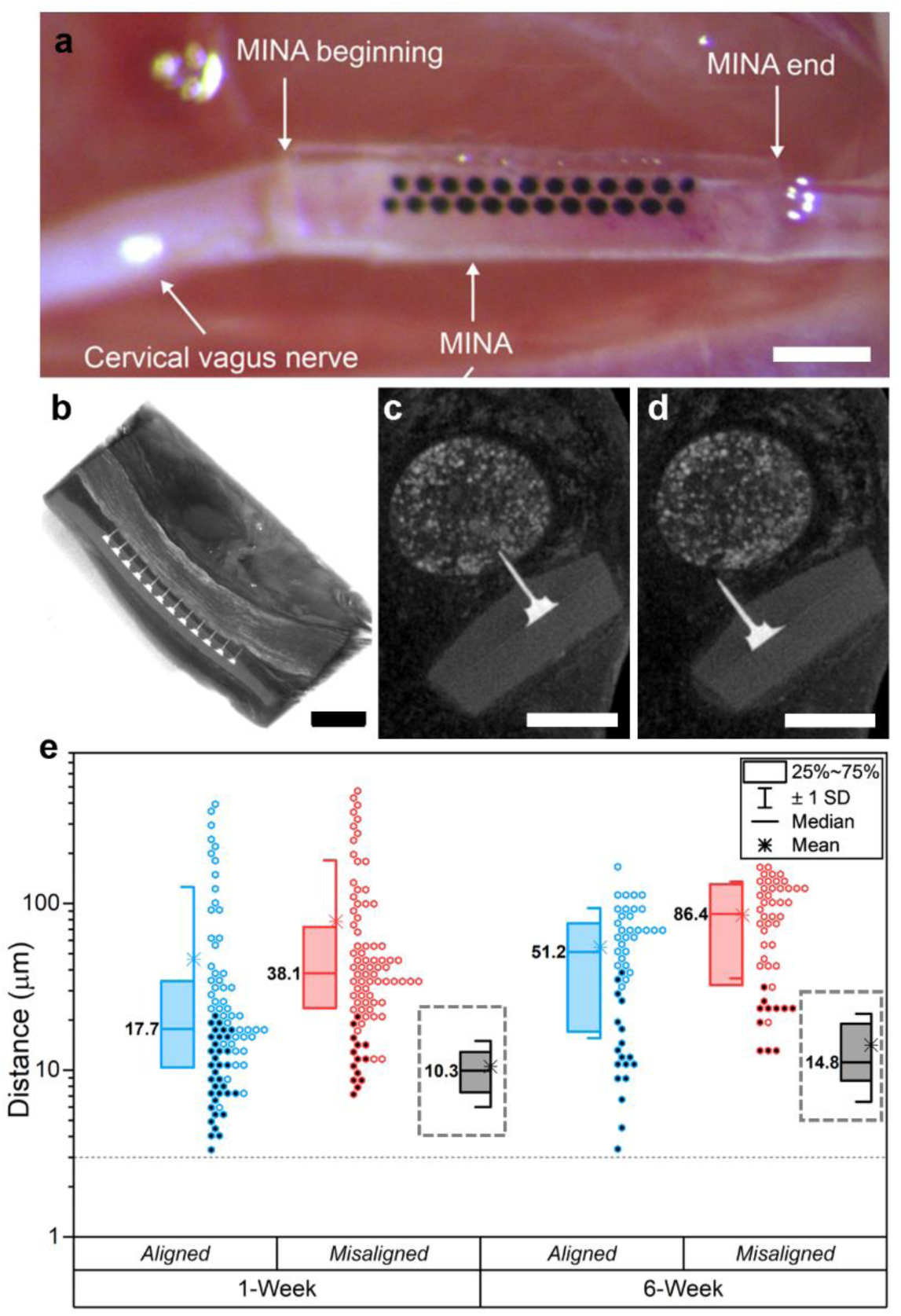
**a)**. Photo of an untethered MINA array attached onto a vagus nerve after laser activated photochemical bonding. **b)**. Side-view image reconstructed from a micro-CT scanning of the device implanted section. Cross-section views of the microneedle electrodes and the implanted nerve. **c)**. One electrode on a well-aligned column and **d)**. A mis-aligned column from the same subject at 1-week timepoint. **e)**. Distance from each electrode to a nearest myelinated fiber. The data points that are colored with black and the grey box plots represent the electrodes inside fascicles. Scale bar in **a-b)** 500 μm; **c-d)** 250 μm.

At the terminal procedure, we observed compound action potential conduction velocities of 1.8-11.0 m/s in 1-week implanted animals and 2.3-10.0 m/s in 6-week animals. These ranges directly overlapped with what we observed in sham animals (1.6-10.6 m/s) and are within the conduction velocity ranges for the myelinated Aδ and B fibers, and unmyelinated C-fibers that are present in the rat cervical vagus nerve^[51,52]^. The stimulation amplitudes to evoke a response were non-significantly higher for 1-week implanted rats (3.8 ± 2.8 mA) versus sham rats (1.5 ± 0.6 mA, p = 0.06), but were unchanged for 6-week rats (2.0 ± 1.1 mA, sham: 2.0 ± 1.0 mA, p = 1.00). The higher stimulation amplitudes at 1 week may have been because the tissue reactivity is near its peak. We observed body weight changes (−6.3 ± 4.0% at 1 week, +22.9 ± 10.6% at 6 weeks) and blood glucose concentration measurements (111.2 ± 15.1 mg/dL at 1 week, 112.0 ± 16.3 mg/dL at 6 weeks) for MINA-implanted rats that were within previously reported ranges for other vagus nerve sham studies^[53,54]^ and were not statistically different from sham animals in this study (p > 0.50 for each comparison).

### 2.5 Micro-CT characterization of nerves with untethered MINA

Micro-CT scanning was used to study the explanted samples as further described in the *Experimental Section*. Micro-CT image stacks when viewed along the axis of the nerve provide a detailed view of spatial orientation of needles and nearby putative axons and changes in tissue density (supporting information videos). The distance between each electrode to its nearest large, myelinated axon was measured in the micro-CT images (Figure 5e). The contrast in our images precludes the identification of small or unmyelinated axons. The nerve section containing MINA was isolated (Figure S5c) and explanted 1 week or 6 weeks after insertion (Figure S5d). Micro-CT imaging of the explant shows that the device remained attached to the nerve. The 1-week micro-CT images show overall closer distance to the fascicles than the 6-week images. Also of note is the axial edges of MINA beyond the microneedles often were curved somewhat away from nerve (Figure 5b). Due to the fine size of the nerve and the challenges of alignment, one of the two columns of the microneedle electrodes was consistently better aligned and inserted than the other column, as shown in Figure 5c and d. Among the 10 untethered MINA that stayed fully attached onto the nerve after 1-week or 6-week implantation, 8 of them had one column noticeably less aligned. After 1-week implantation (N=6, 24 electrodes per implant), the median electrode-to-axon distance of the electrodes on the well-aligned columns was 17.7 μm. After 6-week implantation (N=4, 24 electrodes per implant), the median distance increased to 51.2 μm for the aligned column. 49 out of 144 electrodes, and 30 out of 96 electrodes remained inside the fascicles with thin tissue encapsulation after 1-week and 6-week implantation, respectively. The median thickness of the tissue encapsulation over those electrodes was no greater than 10.3 μm and 14.8 μm after 1-week and 6-week implantation, respectively.

Two processes of the foreign body reaction to the implanted untethered devices were observed from the micro-CT scans. First, the tissue under the device substrate was thickened. We presume that the thickened epineurium and/or adjacent tissue pushed the electrodes out of the fascicles. Second, the part of the electrodes which remained inside the fascicles was encapsulated by a layer of connective tissue. Both processes have contributed to the increase of the electrode-to-axon distance. The mechanical oscillation of the adjacent carotid artery, itself much larger than MINA and the vagus nerve, may have contributed as well. Variance among the initial adhesion strength after photochemical tissue bonding may have also contributed. With imperfect bonding, connective tissues may have grown in between the substrate and the epineurium, which could have resulted in pushing the electrodes out of the fascicles. However, thickening of the epineurium layer is not quantifiable because of the poor contrast between the epineurium and surrounding connective tissue. The sham and MINA-implanted groups at all time points exhibited connective tissue around the nerve and neither our micro-CT nor osmium-tetroxide-stained images provide enough contrast to identify the epineurium boundary.

Previous research on tissue encapsulations over intrafascicular implanted electrodes was studied with a variety of devices including TIME, LIFE, wire electrodes, high-density Utah array, and flexible microneedle electrode array. In one TIME device, the average encapsulation thickness was 135 μm measured after implantation in the median nerves (diameter ∼4 mm) of minipigs for 33 to 38 days^[55]^. In a thorough characterization of a polyimide-based LIFE device implanted in a rat sciatic nerve (diameter 1.0-1.5 mm), the encapsulation area around the device for the period 17 to 165 days was stable at approximately 1.1 mm^2^. Images from this work suggest that at least in some examples, the encapsulation after 3 months was smaller but still at around 125-175 μm^[22]^. For Pt wire electrodes, the encapsulation was around 128 μm after implantation in rodent sciatic nerves for 3 months^[56]^. For high-density Utah arrays, the encapsulation reported was around 540 μm after implanting in human median and ulnar nerves for 28 days^[57]^. In the context of this review of intrafascicular devices, the encapsulation we observed is an improvement for sub-millimeter nerve interfaces.

As a summary, reducing the electrode-to-axon distance in both acute and chronic implantation increases the intrafascicular recording/stimulation quality^[58]^. Our approach succeeded in achieving thinner tissue encapsulation compared with previous research. However, no chronic recordings were attempted and tethering of the nerve cuff would be expected to worsen tissue reactivity. To secure more electrodes inside the fascicles, we recommend increasing the length of the microneedles. Arguably, a simpler manufacturing technique would use aerosol jet printing, which has been demonstrated to achieve a length of more than 1 mm albeit with a blunt tip^[59]^.

### 2.6 Histomorphometry and rehabilitation during 1-week and 6-week implantation

Vagus nerve samples were extracted from 5 groups of animals, including control (N=3), 1-week sham (N=5), 1-week MINA (N=5), 6-week sham (N=3) and 6-week MINA (N=4). Details of the extraction and tissue processing steps can be found in the *Experimental Section*. After tissue processing, nerve sections that were distal and proximal to the implanted regions were sliced, stained with osmium tetroxide, imaged, and stitched with 400X magnification as shown in Figure 6a. Cross-sections of healthy myelinated fibers were counted when distinct donut shaped structures were observed, while potentially degenerated myelinated fibers were identified as solid disks. The entire nerve cross-section at approximately 2-mm proximal and distal to MINA was analyzed in this way. Morphology information including diameter, area, and g-ratio of the myelinated axons were analyzed and collected using Image J (NIH). The average count per subject and the frequency of healthy myelinated fiber with specific diameters are shown in Figure 6 b-e. The means and standard errors of such parameters of myelinated fibers are shown in Figure 6 f-h.

**Figure 6.**
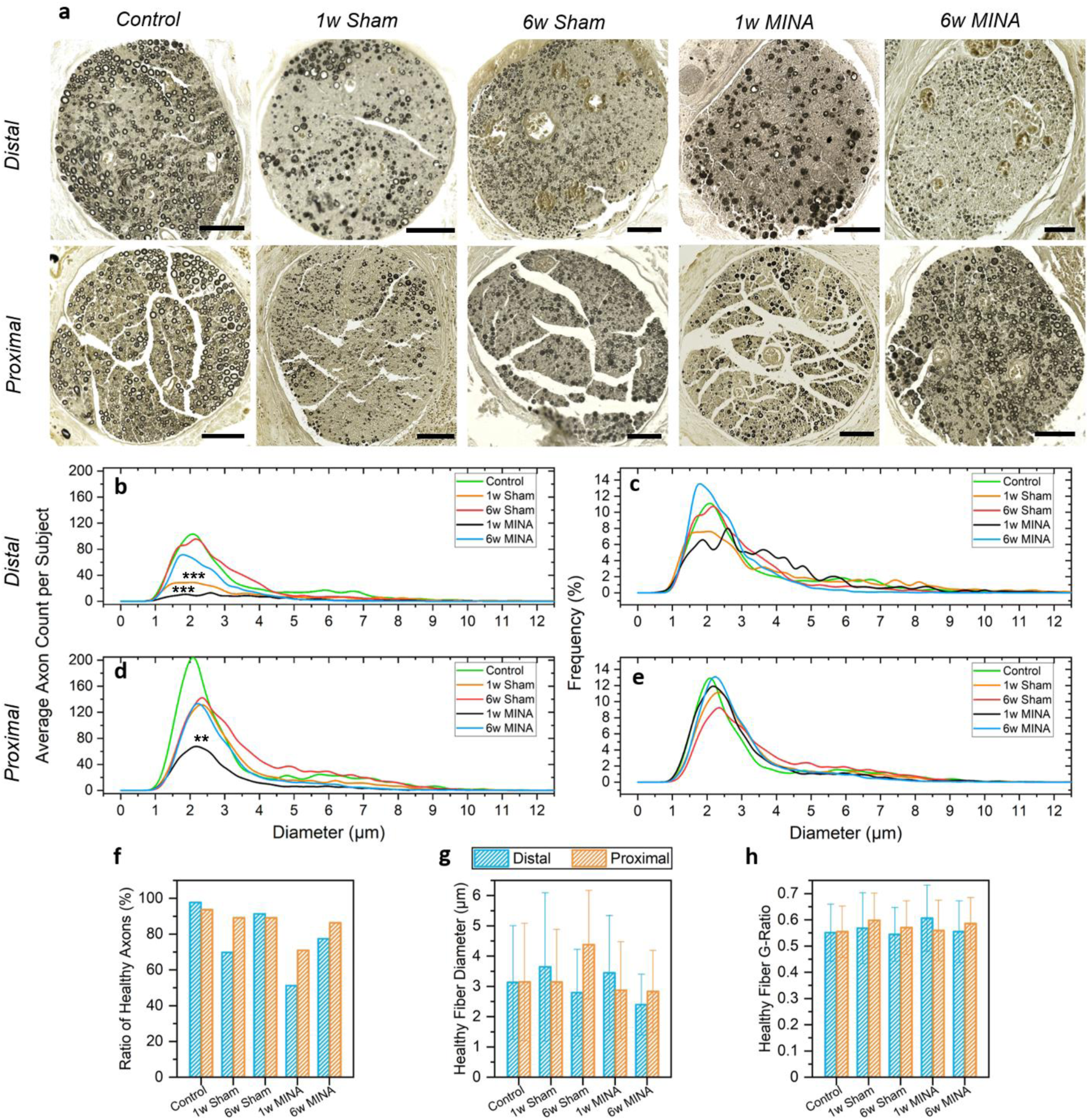
**a)**. Examples of the tissue sections from each group including control (N=3), 1-week sham (N=5), 1-week MINA-implanted (N=5), 6-week sham (N=3) and 6-week MINA-implanted (N=4). All the scale bars are 50 μm. **b)**. to **e)**. Diameter distribution of healthy myelinated axons from each group, average count per subject and frequency (bin size 0.24 μm). 1w MINA was significantly different from the control group when proximal or distal sections were analyzed (** p< 0.01-0.05; *** p<0.001, GLMM, Table S2). 1w Sham was only significantly different on the distal side of the nerve injury. **f)**. Percentage of healthy axons for each group out of the total identified axonal population **g)** and **h)**. Means and standard deviation of the diameters and g-ratios of all the myelinated axons over all the samples from each group.

In all the sham and MINA-implantation trials, nerves were mechanically teased apart from adjacent tissue during the surgery, lifted ∼3 mm upward onto a nerve hook, exposed to green light with a temperature increase of less than 3.5°C, and compressed with a force of approximately 446 ± 187.9 mN to simulate or perform the microneedle electrode insertion (although the nerve itself may have absorbed less force considering misalignment and compression). In additional to compression, a tensile strain due to elevating of the isolated vagus nerve was measured at 5.9±2.5% as described in the previous section.

A generalized linear mixed model was used to examine the statistical difference of the diameter distribution of the healthy fiber diameter between the control group and each other group. Details of the model are described in the *Experimental Section*. Shown in Table S2, we compare all four cohorts with the control group as the intercept for distal, proximal, and combined distal and proximal. The 6-week MINA or 6-week sham groups were not significant in any analysis.

However, the diameter distribution between the control group and the 1-week MINA group was tested to be significantly different (p < 0.001). The results showed that the 1-week sham operation was less healthy than the control nerves (increased abnormal axon count and decreased overall axon count). The results also demonstrated a healthy recovery of the nerve tissues as more time was given post implantation. We believe the sham condition has room for improvement and our results underscore the surgical challenges of implanting microneedle arrays in the cervical vagus nerve in rodents. It also may be possible that implanting MINA in a location further from large pulsations of the carotid artery may have mitigated the observed reactivity.

## 3. Conclusions

In summary, we demonstrate a scalable penetrating electrode array with mechanically independent microneedle electrodes, a micron-sized electrode tip, and a microneedle shaft near the size of a large axon. This device also integrated a thermal oxide insulation layer, which is unique among microfabricated penetrating electrode array. This unique design penetrates just through the epineurium of a nerve and interfaces directly with axons at 1-week, and some electrodes remained close to axons even at 6-weeks. The histomorphology analysis revealed no statistical significance between healthy fiber diameter distribution of the control group and the experimental group after 6-week implantation, although microneedle encapsulation was observed. Our surgery for cervical vagus implantation solved some practical issues with visualizing and attaching a cuff-less array, but also suggested that the surgery itself creates a significant decline in nerve health independent of the presence of a device. The acute recording demonstrated a functioning array and high signal fidelity, which could be used for sensory feedback for closed-loop neuromodulation. Our results also suggest that microneedle electrodes with a 100-μm long structure would be valuable to such a study. With the silicon reactive ion etching approach described in this paper, creating an array with various microneedle electrodes length is also achievable. Integrated optics, such as an endoscope built into the nerve hook, should improve the visibility of the implantation process, which can minimize the nerve manipulation and improve alignment accuracy during electrode insertion. As future work, chronic *in vivo* recordings and stimulations should be performed to further explore the potentials of MINA together with the cuff-less chronic device to tissue bonding approach.

## 4. Experimental Section

### 4.1 MINA fabrication

A simplified fabrication process of MINA is shown in Figure 2. Starting with the fabrication of the silicon microneedle electrode array, we used a 4-inch silicon on insulator (SOI) wafer with a 180 μm thick device layer and a 0.5 μm thick buried oxide (BOX) layer. The device layer was made conductive (ρ = 0.001-0.005 Ω·m) by doping with a high concentration of boron. First, a silicon dioxide layer with thickness of 1 μm was deposited and patterned to form a hard mask for silicon microneedle plasma etching. Then, the device layer was etched with a sequence of anisotropic etching, deep reactive ion etching (DRIE) and isotropic etching. The anisotropic etching at the beginning defined the shape of the microneedle tip. The DRIE process removed the silicon in the field, leaving only the cylindrical microneedle shafts and the ring-shape sacrificial structures used to generate a taper. During this process, a silicon base with diameter of 40 μm and height of 20 μm was formed at the bottom of the microneedle. The last isotropic etching shaped the microneedle shafts into 1 to 2-degree tapered pillars (Figure 3, S1). To create a long-lasting, low defect density and bio-compatible insulation, we oxidize the silicon microneedle in a furnace at 1100 °C for a uniformly 0.5 μm thick thermal oxide layer. The SiO_2_ covered microneedle array was then embedded into a 35 μm thick PDMS (Sylgard-184, 10:1) layer, followed by a 400 nm thick Parylene C diffusion barrier coating. The thermal SiO_2_ and Parylene C layer covering the structure approximately 20 to 30 μm in depth from the tip was later removed via etching. The exposed silicon at this region was then coated with Ti/Pt to create recording electrodes. We bonded the wafer upside down onto a glass carrier wafer using a temporal silicone bonding material (Ecoflex 00-50). Then, the handle and the BOX layer was removed to expose the bottom of the silicon microneedles which were mounted upside down. Another Parylene C layer was coated before patterning metal interconnection traces (Cr/Au/Cr) on the backside. This step produced a wrinkled profile on the PDMS surface which was utilized to create the strain-compliant wrinkled thin film metal interconnection (Figure 3). An additional encapsulation layer of SU-8 (2 μm) was coated and patterned to cover each individual metal trace. Additional metal was sputtered at the metal to silicon contact region and the bonding pads. In the end, an 18-pin Omnetics connector (NPD-18-AA-GS) was connected onto the bonding pads via low temperature solder reflow. After encapsulating the backside with another layer of 35 μm thick PDMS, the MINA was released from the handling glass wafer.

### 4.2 Acute neural recording with MINA on peroneal nerve

All experimental procedures were approved by the University of Michigan Institutional Animal Care and Use Committee (IACUC) in accordance with the National Institute of Health’s guidelines for the care and use of laboratory animals. Non-survival procedures were performed on adult, male Sprague-Dawley rats (Charles Rivers Laboratories International). Anesthesia was induced using 5% isoflurane (Fluriso, VetOne) and maintained using 1-3% isoflurane. Rats were placed on top of a heating pad (ReptiTherm, Zoo Med Laboratories) and breathing rate was monitored every 15 minutes. During the acute recording trials, RB coating and photochemical bonding was not performed. First, the left sciatic nerve was exposed and isolated using gross dissection. Retractors (17009-07, Fine Science Tools) were used to maintain visibility. Then, a microscope (Lynx EVO, Vision Engineering) was used to isolate roughly 2 cm of the distal peroneal nerve branch. The peroneal nerve was lifted and placed on a custom 3D-printed nerve holder (see description below) for insertion of the MINA. The MINA was secured to a vacuum suction adaptor, which was positioned over the peroneal nerve using a 3-axis micromanipulator (KITE-R, World Precision Instruments). The micromanipulator was secured to an optical breadboard (MB1218, Thorlabs) that was underneath the animal. A pen-shaped camera (MS100, Teslong) was positioned to view along the peroneal nerve and perpendicular to the MINA fibers to visualize needle insertion. Immediately prior to insertion, the nerve was rinsed with saline (0.9% NaCl, Baxter International). The vacuum suction adaptor with MINA attached was then gently lowered onto the epineurium to insert the MINA. The ground and reference wires were placed under the skin inside the cavity. Recording was performed using an Intan recording system with a 16-channel amplifier board (RHD2216, Intan Technologies). First, neural recordings were taken during a baseline period without stimuli. Then, cutaneous brushing at the ankle or foot was performed for 10-second intervals between 10-second rest periods to elicit sensory responses in the nerve. A 300 to 1500 Hz digital bandpass filter was applied using MATLAB to remove most of the ambient noise and movement artifact. After filtering, spike detection, sorting, and signal-to-noise ratio calculation was performed using Offline Sorter V3 (Plexon).

### 4.3 Rose bengal coating of MINA

The RB coating process was described in our previous work^[15]^. Briefly, collagen and RB were dissolved separately in 30% ethanol. The collagen-ethanol and RB-ethanol solutions were mixed at a 10:1 ratio. Each MINA surface was treated with oxygen plasma to allow surface bonding prior to application of the RB solution. MINAs were then dried at 50 °C to allow evaporation of the solution and form the RB coating. Individual MINAs were cut with a scalpel blade under a microscope as narrow as possible (∼0.5 by 3.0 mm) with slight extensions on the edges of the needle region to allow handling with fine forceps (Figure S4a). The RB coated MINAs were sterilized at low temperature (37 °C) ethylene oxide in preparation for sterile implantation.

### 4.4 Chronic implantation of the untethered MINA array via photochemical tissue bonding

Untethered MINA device implantation experiments were performed on adult, male Sprague-Dawley rats (0.45-0.64 kg, Charles Rivers Laboratories). All animal procedures were approved by the University of Michigan Institutional Animal Care and Use Committee (IACUC). One day prior to surgery, dexamethasone (0.2 mg/kg) was administered subcutaneously. On the day of the surgery animals were anesthetized with isoflurane (1-5%) and injected subcutaneously with carprofen (5 mg/kg), lidocaine (0.4%), and dexamethasone (0.2 mg/kg). Rats were placed on a heating pad, and temperature and oxygen saturation were measured with a vitals monitor (SurgiVet, Smiths Medical). A midline ventral cervical incision was made to access the left vagus nerve. Under a microscope (Lynx EVO, Vision Engineering), the vagus nerve was isolated (8-10 mm) from the carotid artery and surrounding tissue. Using the microscope camera, an image of the isolated vagus nerve next to a ruler was captured for nerve strain calculations.

We designed a 3D-printed (Form 2, Formlabs) custom nerve holder that secured the nerve in place during insertion and elevated the nerve from fluids and cavity breathing motions. The nerve holder is shown in Figure S5c and further design details are given in Jiman et al. 2020^[60]^. The nerve was placed in the trench of the nerve holder (Figure S4f) prior to MINA insertion. Similarly, a pen-shaped camera was positioned in the cervical opening to visualize the alignment and insertion of MINA needles (Figure S4e, f). The nerve was rinsed with saline, excess fluid was removed with an absorbent triangle (18105-03, Fine Science Tools), and the initial temperature of the nerve was measured with an infrared sensor (IRT0421, Kintrex). The MINA was inserted into the nerve (Figure S4g). The diffusion lens flap on the nerve holder was placed on top of the inserted MINA and the 532-nm laser beam was applied for 360 seconds (Figure S4h). Through the diffusion lens, a total of 360 J/cm^2^ of energy were delivered to the device. The lens flap was removed, and the temperature of the nerve was measured again with the infrared sensor. The process of releasing a MINA-implanted vagus nerve from the nerve holder required extremely accurate handling and could result in applying excess tension on the nerve that led to a MINA detaching. We designed a custom 3D-printed nerve-release tool that would be controlled precisely with a micromanipulator (Figure S4d). After adhesion, the vagus nerve was removed from the nerve holder with the nerve-release tool (Figure S4i) and a second image was captured of the MINA-implanted vagus nerve. The length of the isolated nerve was measured with image analysis software (ImageJ) and the nerve strain was calculated as the percent change in the length from the pre-implant measure. The cervical incision was closed with surgical clips and triple antibiotic topical ointment was applied along the closed incision. A subcutaneous injection of carprofen (5 mg/kg) and dexamethasone (0.02-0.05 mg/kg) were administered daily after surgery for 2-3 days. The health of each animal was checked regularly.

One week after the implant procedure, body weight and blood glucose concentration (via tail prick) were measured, and the surgical clips were removed from the incision under isoflurane anesthesia if the animal was a 6-week implant. Sham animals underwent procedures that were identical to the implantation procedure (including laser application) but without a MINA. No implantation procedures were performed on control animals.

### 4.5 Terminal procedure

One or six weeks after the implant surgery, a terminal procedure was performed under isoflurane anesthesia (1-5%) to assess nerve condition and to extract the vagus nerve and implant. The cervical vagus nerve was accessed similarly to the implant surgery. To assess nerve condition, an electrophysiology test was performed. A stimulation probe (017509, Natus Neuro) was placed on the vagus nerve proximal to the implant region and connected to an isolated pulse generator (Model 2100, A-M Systems) (Figure S5a). A bipolar cuff electrode (0.75 mm inner diameter, 0.5 mm contact spacing, Microprobes for Life Sciences) was placed on the nerve distal to the MINA. Electrical stimulation (1-10 mA, 2 Hz, 200 μs pulse width) was applied to evoke compound action potential neural activity recorded with a data acquisition system (PowerLab, ADInstruments) through the cuff electrode (Figure S5b). The neural recordings were analyzed with MATLAB to determine the lowest stimulation that evoked a response and the conduction velocity for each peak of the response.

### 4.6 Tissue processing of nerve samples

Animals were euthanized with an overdose of sodium pentobarbital (400 mg/kg, IP). A 3-6 mm section of vagus nerve was extracted centered on the implanted section. After extraction, the cervical vagus nerves with implanted MINAs were cut (control nerves were left uncut) into three portions, proximal to the implanted area, the implanted area, and distal to the implanted area, and placed in 3% glutaraldehyde (G5882, Sigma) in deionized (DI) water for 24 hours at 4 °C. Samples were then placed in 0.15 M cacodylic acid (Acros, 214971000) in DI water and stored at 4 °C until ready for staining. All subsequent steps occurred at room temperature unless otherwise noted. To begin the staining process, samples were washed three times in 1x PBS (BP3994, Fisher) with each wash lasting 5 minutes. Next, samples were covered with 2% osmium tetroxide (19152, Electron Microscopy Science) for 2 hours. Samples were then triple washed in DI water, followed by triple washing in 1x PBS with each wash lasting 10 minutes. Finally, samples were quadruple washed in 30%, then 50%, and lastly 70% ethanol in DI water for 12 total washes, with each wash taking 5 minutes. The nerve sample containing the implant was then stored in 1x PBS at 4 °C until micro-CT imaging. Proximal, distal, and control nerve samples were stored in 70% ethanol at 4 °C until paraffin processing.

### 4.7 Micro-CT imaging and analysis

The osmium-stained and device-implanted section of the nerve sample was placed inside a 1.5 mL microcentrifuge tube filled with 1x PBS. The tube was then sealed with a Parafilm sheet to prevent liquid vaporization during the scanning process. Micro-CT imaging was performed using a 3D X-ray microscope (Zeiss Xradia Versa 520). The automatic 3D scanning had a minimum resolution from 1.7 μm x 1.7 μm x 1.7 μm to 3.5 μm x 3.5 μm x 3.5 μm among different trials. 3D intensity data was then reconstructed and visualized by DragonFly (Object Research Systems). The distance between electrodes and the adjacent myelinated axons were annotated manually and calculated automatically with the image analysis tools of DragonFly.

### 4.8 Histomorphometry analysis

Images were taken of the entire nerve cross-section sample using a Keyence BZ-X810 microscope with a 60x oil immersion lens and stitched in the X, Y, and Z directions. Z-focus spanned only 1-μm and improved overall contrast. The parameters measured include: (1) total number of myelinated fibers; (2) axon diameter; (3) fiber diameter; (4) fiber diameter frequency. With the experimenter blinded to the experimental groups, all images were transferred into Adobe Photoshop. Auto-toning correction was made, and a new transparent layer was placed over each entire image. The original image subsequently becomes the background image and is not drawn on. With an X-pen tablet, the entire image was then manually circled or filled in. Clear osmium-stained rings were circled. Solid osmium discs (putative degenerated axons) were filled in completely, and these were counted separately from “healthy” axons. The pressure applied with the pen on the tablet created the thickness needed for the myelin rings. Once the entire 60x image was circled or filled in, the original background layer was removed. The now top layer was then edited so that no rings were touching and fully closed. Once editing was completed, the layer was saved as its own tiff file. This image was then opened in Image J (NIH) and counted. Once opened in Image J each image was changed to an 8-bit image and thresholded. Calibration measurements were inputted into the appropriate fields for these images with the scale being 7.94 pixels/μm. Next, the analyze particle menu is used to find the area and perimeter of objects with and without holes. All these data points were then exported to an Excel template specifically developed by our lab for histomorphometry calculations. Once all the data points were input into the Excel file, the area of the entire nerve image was found by using the polygon tool in Image J. The area of blood vessels and large defects in the nerve samples were measured and subtracted from the total nerve area.

### 4.9 Statistical analysis

Statistical significance for histomorphometry was examined using a generalized linear mixed model (GLMM). The model fitting via maximum likelihood using Laplace approximation was performed in R. Specifically, the fixed and random effect of the linear predictor was the count for each fiber diameter (0.24-μm bins from 0.95 to 12.6 μm) and subject ID, respectively. The axon count is predicted according to a Poisson distribution with its expectation related to the linear predictor given by the mixed effect. The control group was used as the intercept. Standard error and z-score were used to examine the statistical difference between each group and the intercept group. Significance of the statistical difference was annotated based on the z-score value. The body weight, blood glucose concentration, and minimum stimulation amplitudes for each group did not follow a normal distribution (confirmed with Kolmogorov-Smirnov test). A two-sided Wilcoxon rank sum test was used to test for statistical significance in these measures in MATLAB. Statistical significance was considered at p < 0.05. Where relevant, data is presented as mean ± standard deviation.

## Acknowledgements

We thank the Stimulating Peripheral Activity to Relieve Conditions (SPARC) management team, Felicia Qashu and Michael Wolfson in particular, for monthly suggestions to advance this work. DRIE etching and design resulted in the controlled needle shape reported here, for which we thank Saman Parizi, Shuo Huang, Prof. Mark Kushner, Brian van der Elzen, and the Lurie Nanofabrication Facility staff. We thank Hannah Parrish and Zhonghua Ouyang for assistance with nerve sample preparation, and Nancy Senabulya at the Michigan Center for Material Characterization facilitiesfor her technical assistance in micro-CT imaging. We thank Dan Ursu for his advice in the histomorphometry analysis. We thank Lauren Zimmerman, Eric Kennedy, Georgios Mentzelopoulos, Hannah Parrish, Nicolos Buitrago and Nikolas Barrera for their advice and/or assistance with surgical procedures and the Unit for Laboratory Animal Management. We thank Muru Zhou at University of Michigan for her outstanding illustration drawing (Figure 1). This research was supported by the National Institutes of Health (NIH) SPARC Program (Award 1OT2OD024907, 1OT2OD026585) and the Texas Institute for Restorative Neurotechnologies at UTHealth.

