## Supporting Information for "Ultra-flexible and Stretchable Intrafascicular Peripheral Nerve Recording Device with Axon-dimension, Cuff-less Microneedle Electrode Array"

### Support Information

#### Contents

Table S1. Comparison of tensile (t) and shear (s) strengths of various tissue adhesives

Table S2. Fixed-effect analysis using generalized linear mixed model for each Group against Control samples

Figure S1. A variety of silicon microneedle shaping by deep reactive ion etching and reactive ion etching.

Figure S2. Tensile strength measurements of MINA after rose bengal activation.

Figure S3. In vitro temperature measurements during and after rose bengal light activation.

Figure S4. Implantation of rose bengal coated MINA.

Figure S5. Electrophysiology testing and nerve extraction at a terminal procedure.

**Table S1.** Comparison of tensile (t) and shear (s) strengths of various tissue adhesives.

| Adhesive | Strength (kPa) | Substrate | Curing Agent, Time/Dosage | Adhesive Type | Reference & Note |
|---|---|---|---|---|---|
| Rose bengal | 3.41 ± 1.96 (t) | Sciatic nerve, PDMS/ParC | Green light, 300 J/cm$^2$ @ 60W/cm$^2$ | Dry | This Work |
| | 7.12 ± 4.71 (t) | Sciatic nerve, PDMS/ParC (+collagen) | | | |
| | 18.2 ± 13.4 (s) | Sciatic nerve, PDMS/ParC (+collagen) | | | |
| | 34.6 ± 22.6 (t) | Sciatic nerve, PU | | | |
| | 26.7 ± 18.1 (t) | Sciatic nerve, PU (+collagen) | | | |
| Cyanoacrylate | 46.67 ± 12.13 (t) | Porcine vocal folds | Air, 5minutes | Wet | [1] |
| | 68.79 ± 13.29 (s) | Porcine vocal folds | Air, 5 minutes | | |
| | 21 ± 60 (t) | Porcine skin | Air, 60-90 seconds | | [2] |
| | 32.6 ± 89 (s) | Porcine skin | Air, 60-90 seconds | | |
| Fibrin glue | 10.7 ± 6.42 (t) | Porcine vocal folds | Thrombin, 60 minutes | Wet | [1] |
| | 13.86 ± 5.03 (s) | Porcine vocal folds | Thrombin, 60 minutes | | |
| | 0.7 ± 0.6 (t) | Porcine skin | Thrombin, 2 hours | | [2] |
| | 2.2 ± 1.3 (s) | Porcine skin | Thrombin, 2 hours | | |
| UVA-riboflavin [a] | 13.6 ± 1.0 (s) | Cornea | 370 nm, 4.5 min @ 30 mW/cm$^2$ | Wet | [3] |
| Polydopamine | 60.13 ± 7.67 (t) | PDMS-PDA/PDMP | Incubate for 1 hour | | [4] |
| | 28.5 (s) | Adhesive hydrogel to porcine skin | None, Immediate | | [5] |

*Note to Table S1.* Medical-grade polyurethane (PU) as a device surface can offer greater adhesive strength than PDMS, but this elastomer was not available for spin-on or embossing applications.

**Table S2.** Fixed-effect analysis using generalized linear mixed model for each Group against Control group.

| | Group | Estimate | Std. error | z value | Pr (>\|z\|) | |
|---|---|---|---|---|---|---|
| *Distal & Proximal* | *1w MINA* | -1.34673 | 0.28293 | -4.76 | 1.94E-06 | *** |
| | *1w Sham* | -0.46565 | 0.28246 | -1.649 | 0.0992 | . |
| | *6w MINA* | -0.43562 | 0.29565 | -1.473 | 0.1406 | |
| | *6w Sham* | -0.02143 | 0.31577 | -0.068 | 0.9459 | |
| *Distal* | *1w MINA* | -1.793436 | 0.313318 | -5.724 | 1.04e-08 | *** |
| | *1w Sham* | -1.011510 | 0.298228 | -3.392 | 0.000695 | *** |
| | *6w MINA* | -0.534037 | 0.311181 | -1.716 | 0.086133 | . |
| | *6w Sham* | -0.001098 | 0.332431 | -0.003 | 0.997364 | |
| *Proximal* | *1w MINA* | -1.33071 | 0.41129 | -3.235 | 0.00121 | ** |
| | *1w Sham* | -0.27218 | 0.41061 | -0.663 | 0.50741 | |
| | *6w MINA* | -0.386416 | 0.42929 | -0.900 | 0.3680 | |
| | *6w Sham* | -0.08697 | 0.45906 | -0.189 | 0.84973 | |
| | p-value code | <0.001 ‘***’ | 0.001-0.01 ‘**’ | 0.01-0.05 ‘*’ | 0.05-1 ‘.’ | >1 ‘’ |

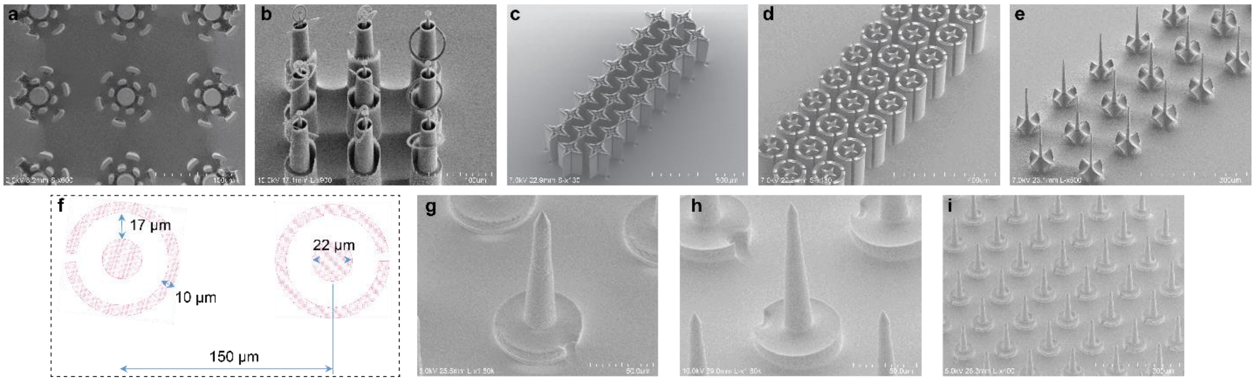


**Figure S1.** A variety of silicon microneedle shaping by deep reactive ion etching and reactive ion etching. **a).** A double pilar arrangement, **b)** double sacrificial shield, **c)** a cross-shaped needle and sacrificial shield, **d)** a cross-shaped needle and circular sacrificial, and **e)** and the narrowest needle attempted, which suffered from fracture during insertion. **f).** Final mask pattern for a 150-µm pitch microneedle. **g).** The post-etch roughness resulted from the scalloping of DRIE and the isotropic plasma etch. **h-i).** After thermal $SiO_2$ was grown on the surface and a brief hydrofluoric acid etch, a smooth surface was formed and was highly scalable.

*Note on Fig. S1:* Our initial attempts at making sacrificial pillars, Fig. S1(a) was inspired by Hanein et al.[6]. This lacked the smoothness and taper control desired for the needle sidewalls, so we settled on sacrificial cylinders with at least one slit for plasma reactant and product exchange. These slits left an artifact structure in the needle base but greatly improved the uniformity of each needle taper. As first described in Yan et al.[7], the timing for each anisotropic and isotropic stage was approximated with modeling of the etch, but ultimately required empirical evaluation to finalize the etch recipe specific to a given tool. Some of the advanced geometries, such as (c) and (d) were not fully optimized for use here but may provide a way to improve the strength of a needle while reducing the cross-sectional area.

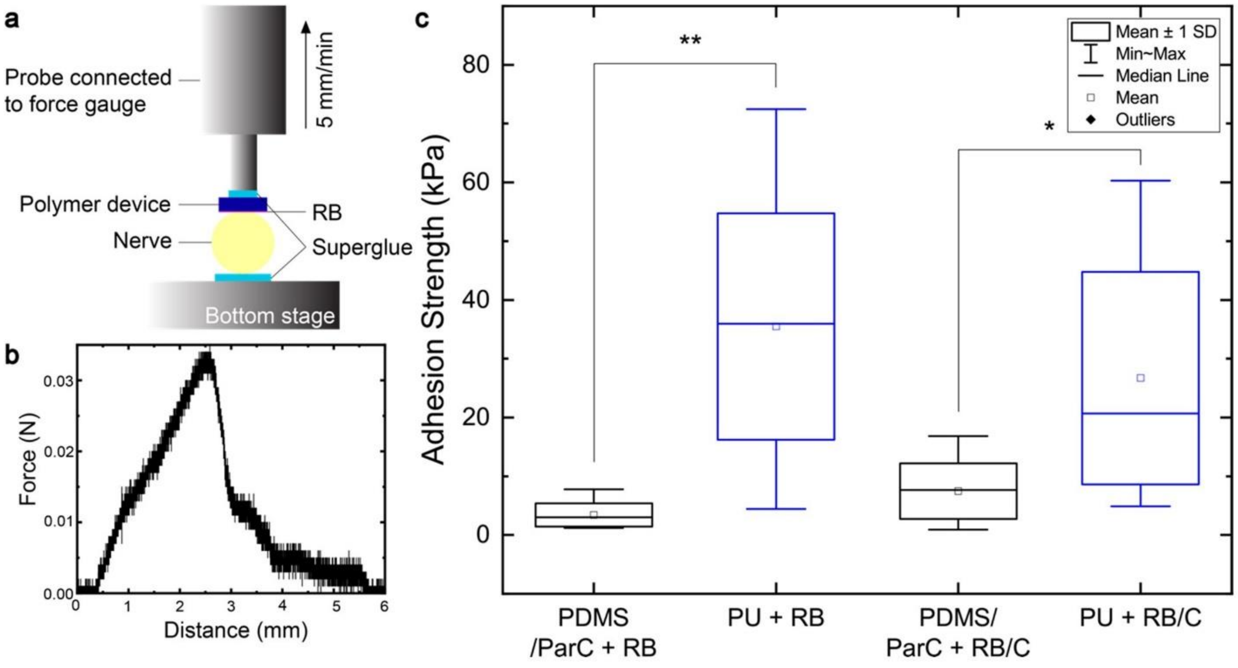


**Figure S2.** Tensile strength measurements of MINA after rose bengal activation. **a).** Tensile adhesion strength testing setup. **b).** Representative pulling force vs. traveling distance during a single tensile adhesion strength test. **c).** Adhesion strength of devices with different surface and coating to sciatic nerves attached via photochemical tissue bonding (N=10).

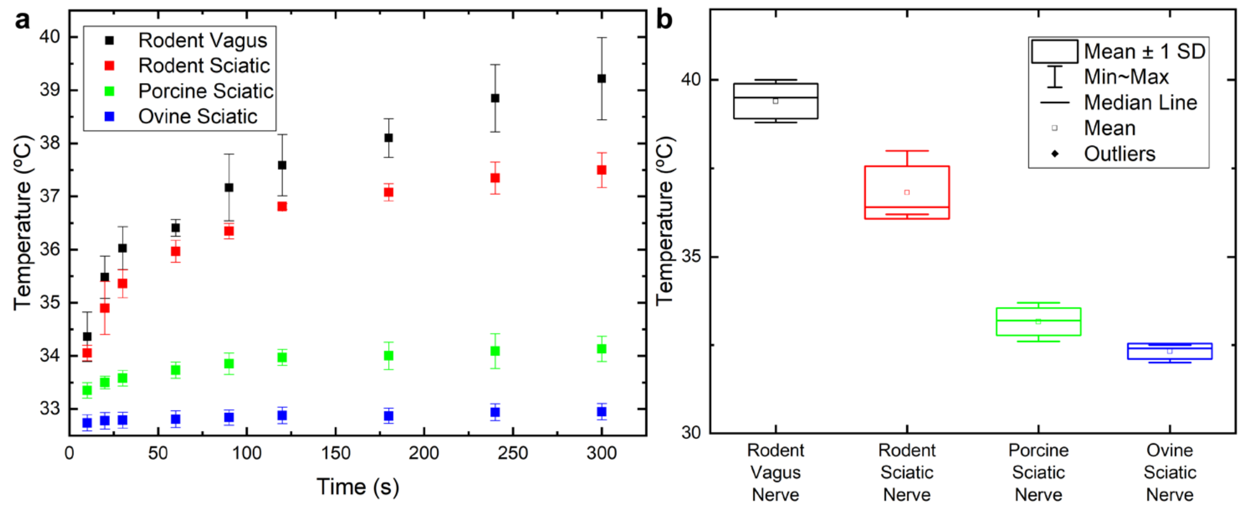


**Figure S3.** *In vitro* temperature measurements during and after rose bengal light activation. **a).** *In vitro* temperature increase of rodent vagus and rodent, porcine, and ovine sciatic nerves upon irradiation. **b).** Maximum temperature increase of rodent vagus and rodent, porcine, and ovine sciatic nerves after 300 seconds of irradiation at 1 W/cm$^2$ (N=5).

*Note on Fig. S3:* The nerve samples used here were extracted and kept refrigerated in saline for less than 24 hours. Before measurement, the saline was heated to 37 °C. The samples were loaded onto the trench of the lens-attached nerve holder (Figure S4c). Irradiation was applied through the diffusion lens. Temperature was measured with an infrared camera (Seek Thermal®)

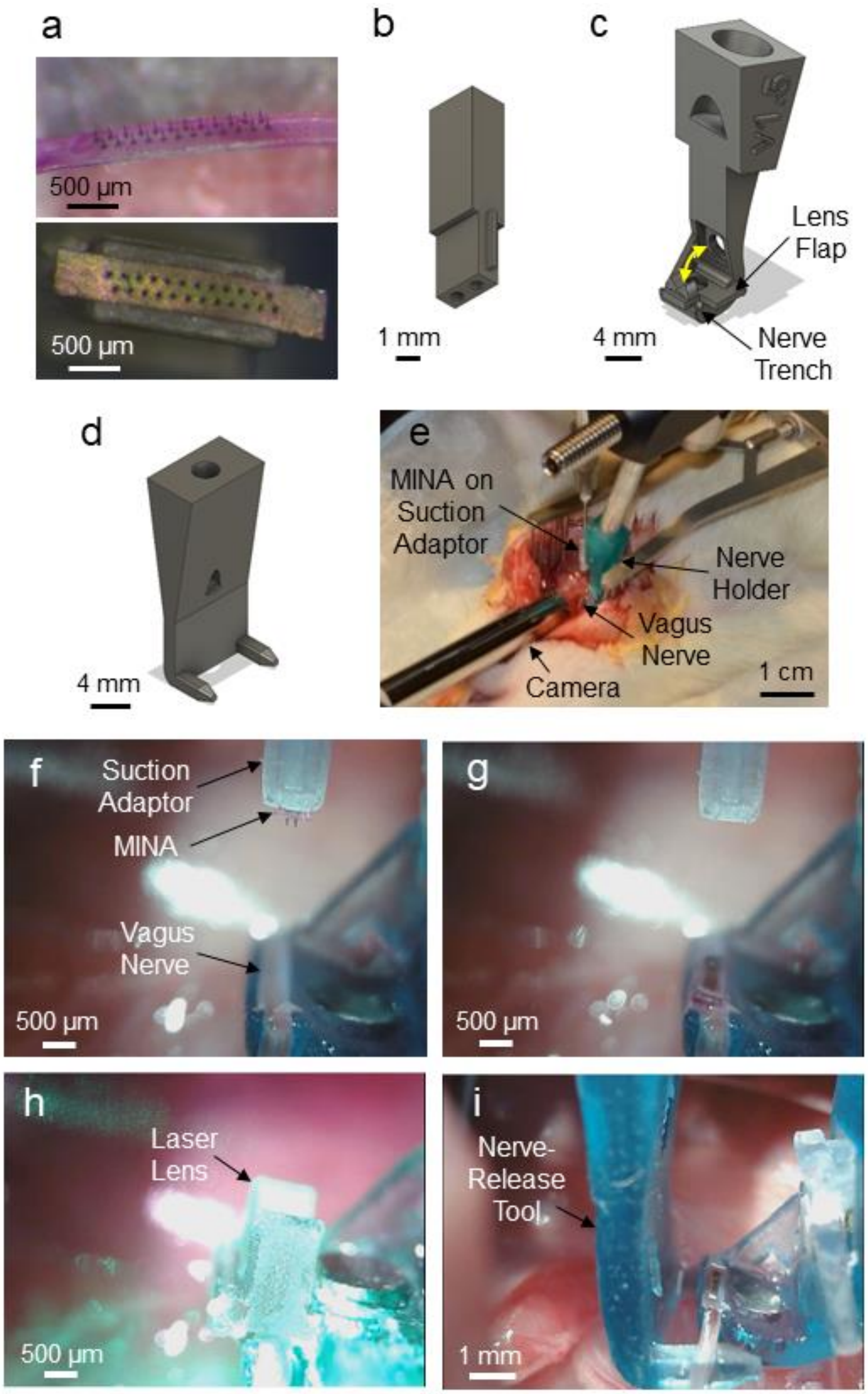


**Figure S4.** Implantation of rose bengal coated MINA. **a).** Rose bengal coated MINA with 2x12 needle configuration (top). MINA centered on vacuum suction adapter (bottom). **b).** Design of vacuum suction adapter. **c).** Nerve-holder design. Yellow arrows show movement path of lens flap. **d).** Design of a separate nerve-release tool. **e).** Surgical setup for implantation in vagus nerve. **f).** View from pen camera for aligning MINA. **g).** MINA inserted in vagus nerve. **h).** Placement of lens flap and activation of rose bengal coating with laser. **i).** Release of MINA-implanted vagus nerve from the nerve holder with the nerve-release tool.

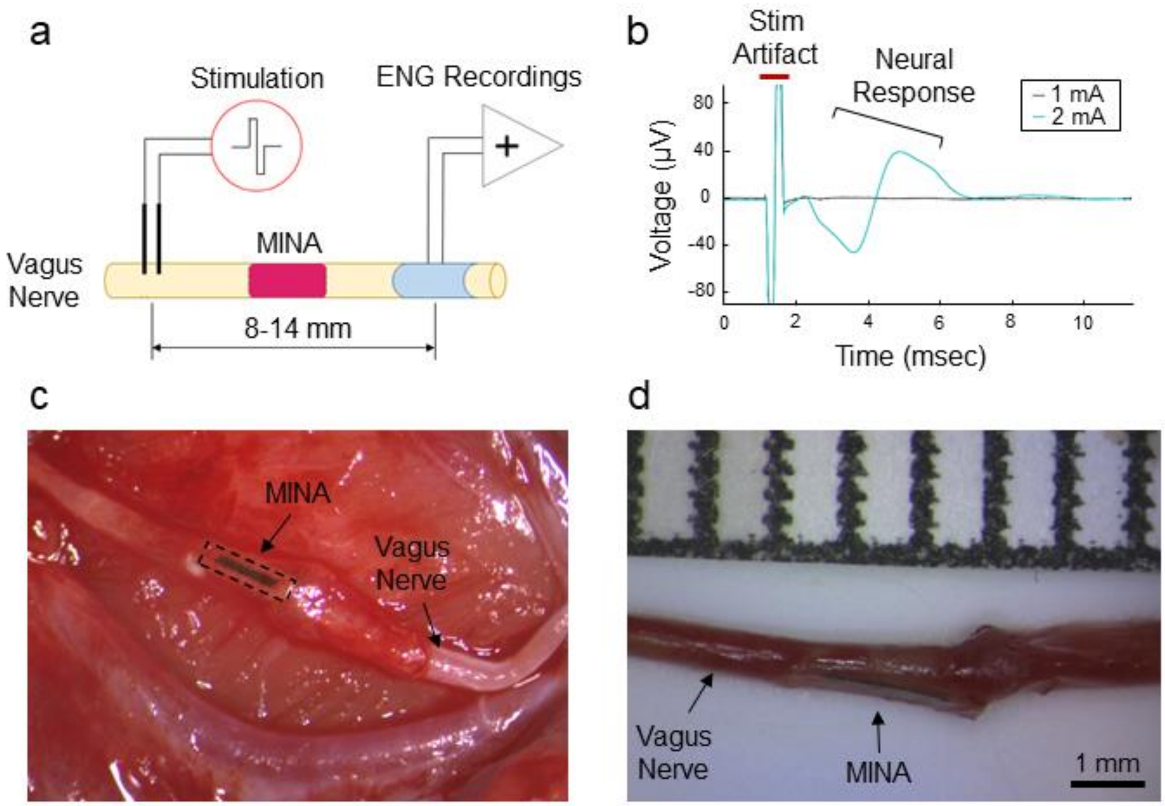


**Figure S5.** Electrophysiology testing and nerve extraction at a terminal procedure. **a).** Setup diagram for electrophysiology testing. **b).** Example stimulation-evoked compound action potential responses in one rat. **c).** Isolation (top-down view) and **d).** extraction (side view) of MINA-implanted vagus nerve for a 1-week rat.
